# Developmentally regulated generation of a systemic signal for long-lasting defence priming in tomato

**DOI:** 10.1101/2023.10.09.561512

**Authors:** Katie Stevens, Michael R Roberts, Katie Jeynes-Cupper, Marco Catoni, Estrella Luna

## Abstract

Tomato plants can be chemically primed to express long-lasting induced resistance (IR) for the protection of fruit against pathogen infection. Here, we determined that priming results in maintenance of IR in fruit and progeny of tomato plants only when initiated at an early developmental stage. Global DNA methylation analysis revealed seedling-specific changes, which occurred in the context of lower basal methylation, suggesting a greater epigenetic imprinting capacity of young plants. Remarkably, IR was found to be transmissible from primed rootstock to grafted unprimed scions. In these scions, we identified a subset of mobile 24 nt small RNAs associated with genes with enhanced expression in response to *Botrytis cinerea* infection in fruit, suggesting the functional association of a systemic signal with long-lasting IR and priming. Through integrated omics approaches we have identified markers of long-lasting priming in tomato fruit which could also serve as targets for durable resistance in other crops.

## Introduction

The current food supply chain experiences major losses at the postharvest level due to both injury and infection by pathogenic fungi^1,2^. Tomato is a major global commodity, with 182.3 million tons of fruit produced in 2019^3^. However, its yield is heavily restricted due to pathogens, and 50% of yield loss occurs at the postharvest stage^3^. Postharvest pesticide use is not permitted for tomato fruit in commercial settings^4^, and the main control measures at this stage are limited to cold temperature storage and strict hygiene measures^5^. However, postharvest pathogens such as *Botrytis cinerea*, the causal agent of grey mould, cannot be successfully controlled with these strategies. *B. cinerea* affects plants and fruit of 230 crop species and its infection postharvest causes economic losses of up to $100 billion^6^. Therefore, new strategies are required. A better understanding of tomato defence mechanisms would allow researchers to design strategies to control pre- and postharvest fungal infections and reduce yield waste.

The ‘adaptive’ component of the plant immune system can be referred to as priming^7^. Unlike direct activation of defence mechanisms, which induces significant metabolic alterations, priming minimises energetic costs via targeted allocation of energy resources upon attack, thus resulting in a faster and stronger activation of defence mechanisms when required^8^. Priming is considered to be broad spectrum and has been described in many different plant species, from *Arabidopsis thaliana* to *Malus pumila* (apple trees)^9–13^. Importantly, priming has been shown to be long-lasting^14–17^ and to be transmitted to following generations^18–20^. A very well characterised priming chemical is the non-protein amino acid β-aminobutyric acid (BABA), first identified in the 1960s^21^. BABA has subsequently been documented to be effective against both abiotic and biotic stresses in a range of species^10^. BABA induced resistance (BABA-IR) is associated with a range of changes to the plant such as enhanced physical protection through callose deposition, PATHOGENESIS-RELATED1 (PR1) protein accumulation and increases in defence hormones such as salicylic acid (SA) and jasmonic acid (JA)^13,22–25^. In Arabidopsis, BABA binds to an aspartyl-tRNA synthetase^26^ and changes the canonical function of the enzyme into priming. In tomato and Arabidopsis, BABA can be absorbed through the roots and is then translocated to aerial tissue^16,27^. Although the receptor has not been identified in tomato, BABA is thought to work in a similar way in tomato to Arabidopsis^26^, leading to durable enhanced resistance against *B. cinerea*^28^. BABA treatment has been shown to lead to long-lasting protection of fruit tissue when applied at the seedling stage, thus conferring postharvest protection^16,29^. Therefore, long-lasting priming offers an alternative approach to fungicides towards protecting plants from postharvest pathogenic infections.

Long-lasting priming has been linked to epigenetic changes such as DNA methylation and the production of small RNAs (sRNAs), as they can contribute to changes in gene expression^14,19,20,30,31^. For instance, analysis of Arabidopsis epigenetic recombinant inbred lines (epiRIL) demonstrated that hypomethylated loci enhanced priming of SA-dependent and SA-independent defences against virulent *Hyaloperonospora arabidopsidis*^32^. Moreover, sRNAs produced by the plant specific RNA-directed DNA methylation (RdDM) pathway have been associated with long-lasting and transgenerational induced resistance in Arabidopsis^19^. Recent work has illustrated that JA-IR is regulated by DNA-demethylation pathways, requiring an intact sRNA binding protein AGO1 to prime defence associated genes^33^. BABA has also been shown to be associated with important changes in DNA methylation. In tomato, global changes to DNA methylation in the CHH cytosine context (H indicates any nucleotide other than G) have been associated with long-lasting BABA-IR in the Money-Maker cultivar. Whilst many differentially methylated regions (DMRs) were found in promoters of differentially expressed genes during *B. cinerea* infection, the majority of primed genes were not differentially methylated^14^. Therefore, the mechanisms behind the long-lasting epigenetic nature of priming are still unclear. In addition, the long-lasting nature of BABA-IR has yet to be explored and utilised for its potential role in postharvest research. Interestingly, tomato plants have been shown to have different methylation profiles depending on both fruit developmental stage and tissue type: CG and CHG methylation levels are lower in fruit tissue than 4-week-old leaf tissue, with the reverse pattern seen in CHH context^34^. However, how changes in developmental stage-dependent DNA methylation mediate the imprinting and the maintenance for long-lasting postharvest priming is unexplored.

Here, we found that the plant’s developmental stage has a major influence on the ability to establish long-lasting priming against *B. cinerea*. We assessed the impact of BABA treatments on a transcriptomic and epigenomic level at different developmental stages and used methylome analysis to test the hypothesis that young plants display greater epigenetic plasticity. Additionally, we found that long-lasting BABA-IR is transmissible to naive scion tissue when grafted on primed rootstock, and we investigated the association of sRNAs with resistance. Through the integration of omics analyses we have identified markers associated with long lasting BABA-IR in tomato for the control of *B. cinerea* in fruit postharvest.

## Results

### BABA-induced resistance occurs in tomato if plants are treated at an early developmental stage

A time course experiment was designed to investigate the dynamics of BABA-IR establishment in tomato (Fig 1a). We used a soil drench approach to prime with BABA the roots of plants at two different developmental stages, either at 2 or 12 weeks after germination (hereafter referred to as BABA2 and BABA12, respectively). We analysed plant material at three developmental times, including leaves collected at 3 and 13 weeks post germination (referred to as T1 and T2, respectively) and in detached ripe, red fruit (referred to as T3). Mock-treated plants were used as controls for each of the timepoints considered.

**Figure 1:**
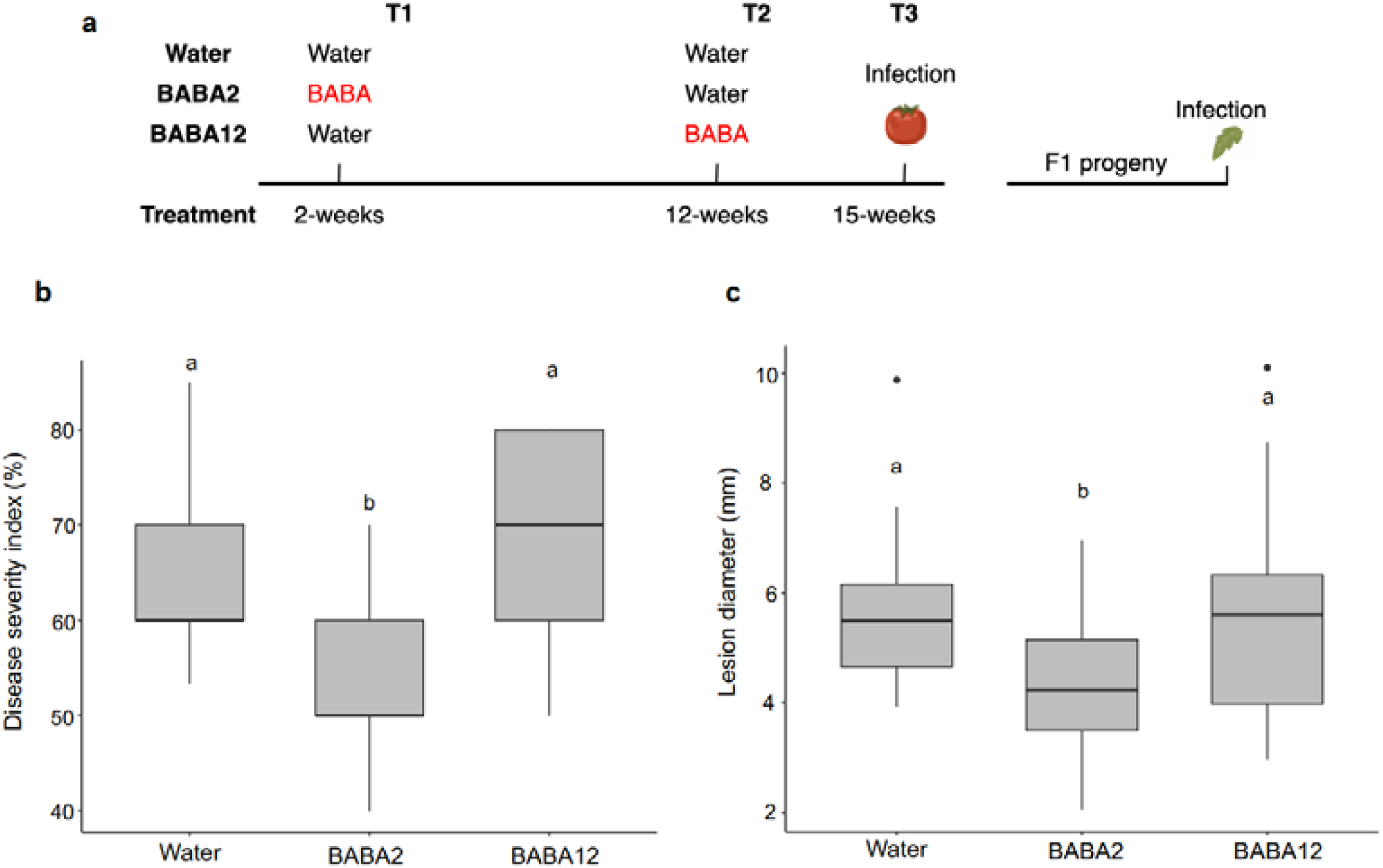
Phenotype data a) Experimental setup. Plants were treated with 0.5 mM BABA via soil drench at either 2 weeks (BABA2) or 12 weeks (BABA12). Resistance against *Botrytis cinerea* was tested in red ripe fruit tissue (collected at 15 weeks) and in leaf tissue from the F1 progeny. b) Disease severity index in ripe tomato fruit converted from the percentage of lesions in 6 disease categories at 2 days post infection (dpi) with *B. cinerea.* c) Lesion diameter in 3-week-old tomato leaves caused by *B. cinerea* at 2 dpi. Different letters denote significant differences among treatment groups (one-way ANOVA, Tukey post hoc test, *p* < 0.05, n = 8-10).

We assessed *B. cinerea* disease progression based on the size of lesions appearing after infection following inoculation of detached fruit harvested from treated plants, and in leaf tissue of the following plant generation. We observed long-lasting resistance only in fruit collected from BABA2 plants and transgenerational resistance in their progenies (*p <* 0.05), while the infection of fruits from BABA12 plants and their progeny did not differ from that observed in mock-treated plants (Fig 1b and c). Although BABA can be phytotoxic in young seedlings, BABA2 treatment caused no significant reduction of relative growth rate (RGR) (Supp Fig 1).

### Short-term and long-term BABA priming of gene expression

To test the priming mechanisms activated by BABA over the short- and long-term, transcriptomic analysis was conducted on *B. cinerea* infected leaf (T1) and fruit tissue (T3). We analysed samples when disease symptoms were first observed at 24 and 48 hours after infection in leaves and fruit, respectively. Principal component analysis (PCA) revealed a clear separation between mock and infected samples in the short-term response. Furthermore, PC1 also differentiated BABA2 from control infected leaves (Fig 2a). In fruit tissue the impact of the infection was more pronounced (Fig 2b), but all the infected samples, regardless of BABA treatment, clustered together. This result indicates that the BABA treatment has a stronger effect on gene expression in leaves shortly after its application, than in fruit at T3.

**Figure 2:**
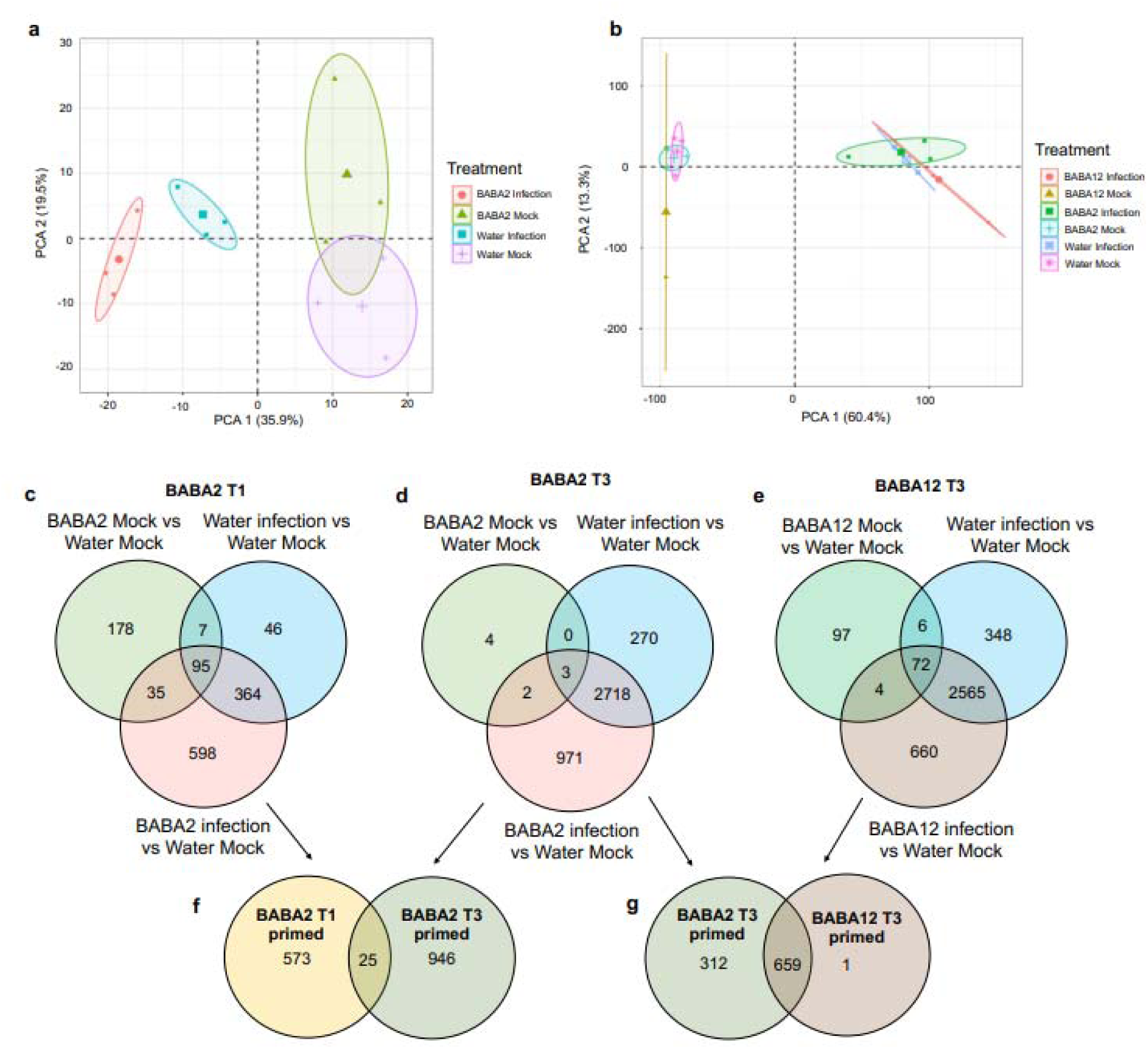
Transcriptome responses to *Botrytis cinerea* infection. Principal component analysis (PCA) of transcriptomic responses to *Botrytis cinerea* infection or mock inoculation in a) early BABA treated (BABA2) and control (water) 3-week-old tomato leaf tissue and b) BABA2, water and late BABA treated (BABA12) fruit tissue. Venn diagram comparisons between treatment-specific differentially expressed genes: c) BABA2 mock vs water mock, water infection vs water mock and BABA2 infection vs water mock at T1, d) BABA2 mock vs water mock, water infection vs water mock and BABA2 infection vs water mock at T3, and e) BABA12 mock vs water mock, water infection vs water mock and BABA12 infection vs water mock. f) BABA2 T1 primed genes vs BABA2 T3 primed genes, g) BABA2 T3 primed genes vs BABA12 T3 primed genes.

To identify genes that were common to responses at T1 and T3, and those that were unique to a single timepoint, comparisons were made between treatment-specific differentially expressed genes (DEGs; Fig 2c-g). Considering that we observed priming-based induced resistance in fruit of BABA2 plants but not BABA12 and mock treated samples, we hypothesised that by comparing the expression profiles among these conditions during infection, we could distinguish molecular determinants of BABA-IR from other unrelated effects of the BABA treatment. Primed genes were identified by filtering out those regulated by BABA treatment alone and those regulated by infection in the absence of BABA (Fig 2c-e). These genes were labelled as ‘BABA2 T1 primed’ and ‘BABA2 T3 primed’, depending on the timepoint when they were identified. A total of 598 and 971 differentially regulated genes were isolated in ‘BABA2 T1 primed’ and ‘BABA2 T3 primed’, respectively. Therefore, it appears that early BABA treatment impacts a greater number of primed genes over the long-term compared to the short-term. We also found 660 differentially expressed genes unique to BABA12 infection at T3 (labelled ‘BABA12 T3 primed’). To assess how similar the differentially expressed genes after infection are between T1 and T3, ’BABA2 T1 primed’ and ‘BABA2 T3 primed’ genes were compared (Fig 2f). Only 25 genes out of a total 1569 genes were shared, indicating that both the timing of the expression of priming and the plant tissue are major drivers of the transcriptomic priming profiles. Finally, comparing ‘BABA2 T3 primed’ genes with ‘BABA12 T3 primed’ indicated that 659 genes of a total 1,631 were shared (Fig 2g). However, since BABA12 displayed a susceptible phenotype (Fig 1b and 1c), it is likely that those shared genes are not directly related to the enhanced resistance phenotype. Therefore, only the 312 genes found differentially expressed exclusively in the early BABA2 treatment, which could be linked to long-term resistance, were considered in subsequent analyses.

### DNA methylation changes induced by BABA are dynamic and increase over time

To explore whether epigenetic mechanisms are involved in long-term priming, we investigated changes in DNA methylation following treatments with BABA in each of the samples collected at the different timepoints of our analysis. We conducted whole genome bisulfite sequencing (WGBS) to obtain DNA methylation profiles of each sample at single cytosine resolution. No significant response to BABA treatment was seen at the global methylation level when each treated sample was compared to the corresponding mock control at the same timepoint (Fig 3a-c). However, we observed a substantial variation in global DNA methylation depending on tissue type and developmental stage. For CG, we observed lower levels of global methylation at T3 (fruit; 77.9%) than at T1 and T2 (leaves; 82.9% and 83.6%) (Fig 3a). Changes associated with the developmental stage were also observed in the CHH context, where global levels increased from 7.3% to 13.5% between T1 and T2 and to 15.4% in T3 in all treatment groups (Fig 3c). However, no variation in CHG methylation levels were found for either tissue type or developmental stage (Fig 3b).

**Figure 3:**
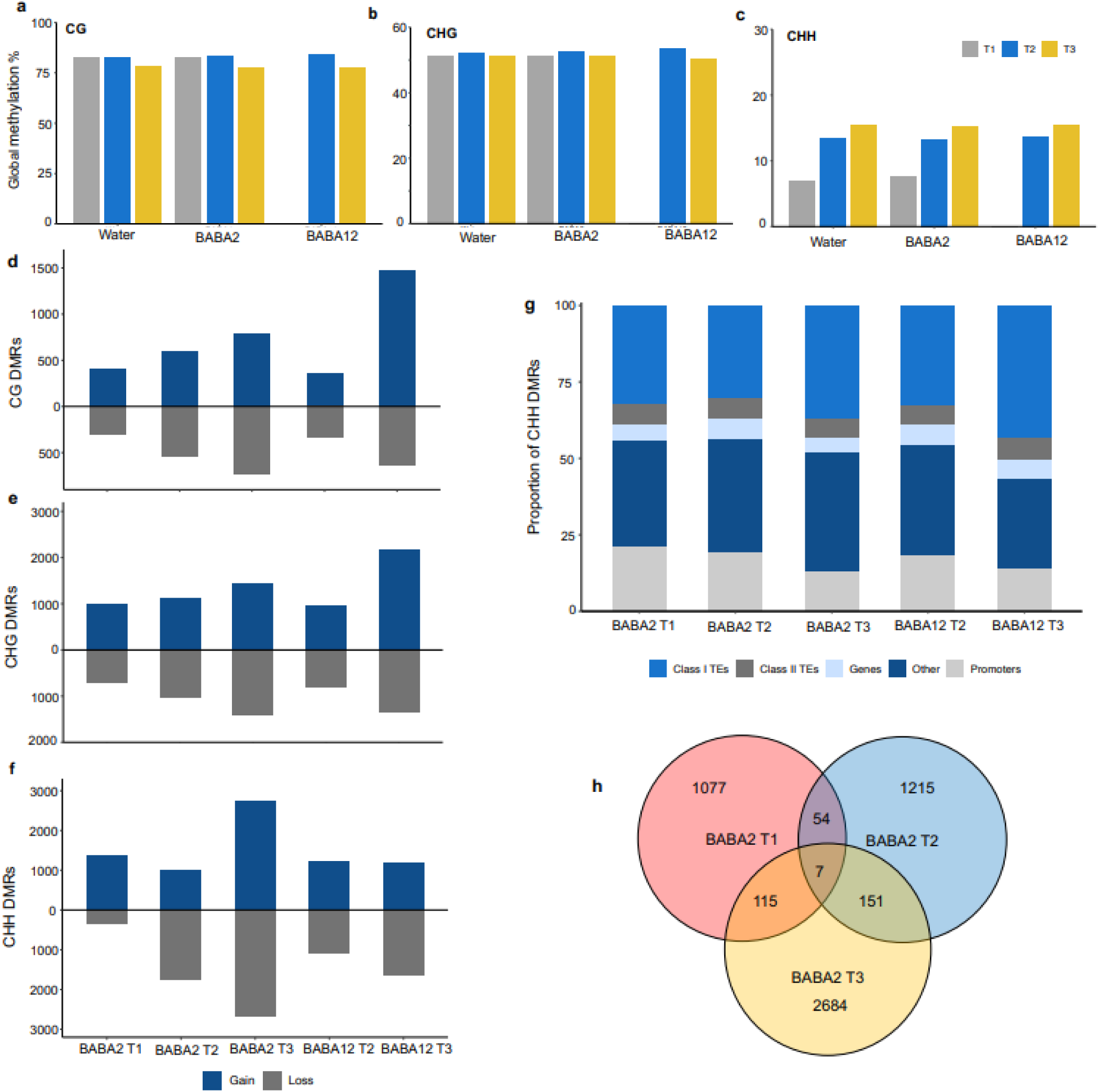
WGBS analysis. Global methylation level across treatment groups in 3-week-old leaf tissue (T1; grey), 13-week-old leaf tissue (T2; blue), and ripe fruit tissue (T3; red) in each cytosine context: a) CG, b) CHG, and c) CHH. Total number of hyper-DMRs (blue) and hypo-DMRs (grey) across samples in each cytosine context: d) CG, e) CHG, f) CHH. g) Location of DMRs across genomic figures in BABA2 leaf tissue at 3-weeks. h) Venn diagram illustrating correspondence between genes overlapping DMRs unique to BABA2 treated plants at T1 (red) T2 (blue) and T3 (yellow).

To investigate the presence of specific DNA methylation patterns associated with BABA priming, we identified differentially methylated regions (DMRs) from samples treated with BABA and the corresponding mock control, for each timepoint. The total numbers of DMRs found in each context are listed in Table 1.

**Table 1:** Total number of 200 bp DMRs* in each cytosine context (CG, CHG, CHH). DMRs* compared to corresponding water control for each treatment group at each timepoint.

| Timepoint | Treatment vs | CG | CHG | CHH |
| --- | --- | --- | --- | --- |

|  | **water** |  |  |  |
| --- | --- | --- | --- | --- |
| **T1** | BABA2 | 714 | 1,720 | 1,751 |
| **T2** | BABA2 | 1,125 | 2,137 | 2,777 |
|  | BABA12 | 692 | 1,783 | 2,297 |
| **T3** | BABA2 | 1,518 | 2,858 | 5,425 |
|  | BABA12 | 2,104 | 3,552 | 2,836 |

\*Differentially methylated regions

Generally, the number of DMRs increases over time in all three cytosine contexts. This effect is most dramatic in the CHH context where the total number of DMRs increases over 3 times from T1 to T3 and 2 times from T2 to T3 in BABA2 plants (Table 1). In contrast, there was only a small increase in CHH DMRs between T2 and T3 for BABA12. There was a more noticeable increase in the numbers of CG and CHG DMRs in BABA12, which was not as pronounced in BABA2. Collectively, these results suggest that DNA methylation in tomato is primarily affected by the developmental stage, displaying increased variation in the CHH context and, less markedly, in the CG context. These results also indicate that the most prominent effect of BABA treatment occurs several weeks after the treatments, with the maximum difference measured in fruits, and many new DMRs arising weeks after BABA2 treatment.

Major differences were identified in the gain and loss of the methylation status of DMRs depending on the timepoint of BABA application, particularly in the CHH context (Fig 3d-f). At T1 BABA2, DMRs were more often hyper- than hypo-methylated, with the greatest effect observed in the CHH methylation context, where 4 times more DMRs were hypermethylated. This gain of CHH DMRs was not maintained in the later timepoints: CHH DMRs at T2 showed greater loss and at T3 gain and loss was equal (Fig 3f). In contrast, BABA12 did not lead to the same pattern of DMR imbalance in any cytosine context and gain/loss ratios were maintained at T2 (Fig 3d-f). At T3 however, there was a gain of methylation in CG and CHG contexts, and a loss in CHH methylation. The annotated features on the tomato genome were used to assess the overlap of BABA2 and BABA12 associated DMRs with features of interest. We observed that for all samples analysed, DMRs were primarily concentrated on Class I TEs for all cytosine contexts (Fig 3g), but also overlap a portion of genes and/or their promoters. Therefore, to identify differentially methylated genes (DMGs) that potentially contribute to priming, we isolated DMGs unique to the BABA2 phenotype. We identified 1,253 DMGs unique to BABA2 at T1, 1,427 at T2, and 2,957 at T3. Overlap of these DMGs revealed that only 54 were present in both T1 and T2, 115 in T1 and T3, 151 in T2 and T3. Only 7 DMGs were present at all three timepoints (Fig 3h). This indicates that early DMRs generated by BABA soil drench treatment at 2-weeks are generally not maintained into older leaves or fruit tissue.

### Genes associated with priming are not frequently methylated

To test whether transcriptional responses associated with BABA treatments are epigenetically regulated, DEGs resulting from the BABA treatment alone (i.e., BABA mock - without infection; Fig 2c-e) and ‘BABA2 T1 primed’, ‘BABA2 T3 primed’ and ‘BABA12 T3 primed’ genes (Fig 2f and g) were compared with DMGs. Venn diagram analysis revealed that very few genes associated with a BABA transcriptional response were also differentially methylated (Fig 4a-c). At T1, only 25 out of 573 ‘BABA2 T1 primed’ genes were also differentially methylated in any cytosine context, the majority of which were hypermethylated and in the CHH context (21/25 and 13/25, respectively) (Supp Table 1; Fig 4a). At T3, no DEGs responding to early BABA treatment without infection were differentially methylated and only 16 genes were both ‘BABA2 T3 primed’ and differentially methylated (Fig 4b). Similarly to timepoint T1, a greater number of DMGs were in the CHH context but there was a more even distribution between methylation status and gene expression (9 hypermethylated DMRs, 9 down-regulated genes) (Supp Table 1). Whilst the data presented in Fig 2 and Fig 3h show that the effects of BABA on gene expression and DNA methylation vary substantially across the different treatment and sampling points, we wondered whether there might be greater similarity in biological impacts when viewed at higher levels of organisation. We therefore applied gene ontology (GO) term enrichment analysis to the full non-redundant lists of DEGs and DMGs contained in the Venn diagrams in Fig 4a-c. However, there was very little similarity between the biological processes impacted by BABA2 at T1 (Fig 4d), BABA2 at T3 (Fig 4e) and BABA12 at T3 (Fig 4f). This reinforces the conclusion that different pathways are associated with short-term and long-term priming mechanisms generated by BABA and that the timing of application is crucial for the priming of specific pathways that result in enhanced resistance.

**Figure 4:**
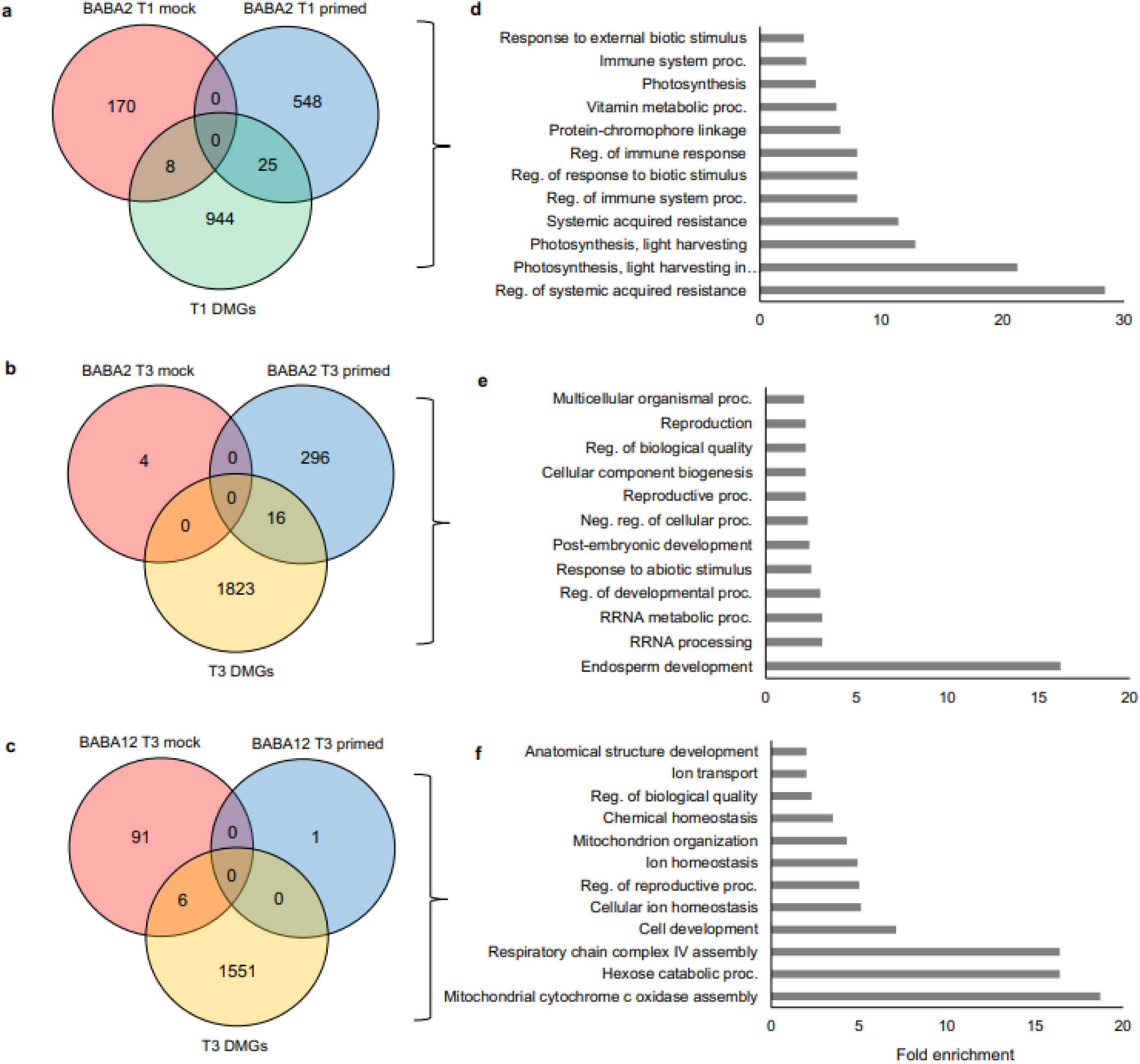
Overlap of all BABA DEGs and genes containing DMRs, and their enrichment. a) BABA2 T1 genes (red representing BABA2 mock DEGs, blue representing BABA2 T1 primed DEGs, green representing T1 BABA2 DMGs), b) BABA2 T3 genes (red representing BABA2 mock DEGs, blue representing BABA2 T3 primed DEGs, yellow representing T3 BABA2 DMGs), c) BABA12 T3 genes (red representing BABA12 mock DEGs, blue representing BABA12 T3 primed DEGs, yellow representing T3 BABA12 DMGs), d) GO term enrichment of all BABA2 T1 genes, e) GO term enrichment of all BABA2 T3 genes, f) GO term enrichment of all BABA12 T3 genes.

### Long-lasting resistance is transmitted from BABA-primed rootstocks to scions across a graft junction

We observed that BABA2 treatment triggers long-lasting IR in tissue which develops after the treatment, and that DMRs specifically associated with BABA2 emerge weeks after the treatment. These observations suggest that long-lasting resistance after BABA treatment may be maintained via a systemic signal. To test this hypothesis directly, grafting experiments were conducted to determine whether BABA-treated tissues could transmit resistance to untreated tissues. We grafted naive scions obtained from plants mock-treated with water onto BABA2-treated rootstocks (hetero graft; HG). As controls, we generated homografts made using both rootstocks and scions obtained from plants treated with BABA (BABA graft; BG) or mock-treated (mock graft; MG) (Fig 5a). Resistance against *B. cinerea* was tested in scion leaf tissue 6 weeks after grafting (corresponding to 6 weeks and 3 days post BABA treatment) and in ripe red fruit tissue. Remarkably, we observed that scion and fruit from HGs displayed the same level of resistance as observed in BGs, with significantly reduced disease symptoms compared to the MG condition (Fig 5b-c). Therefore, our results indicate that BABA-IR is transmissible to untreated scions by grafting.

**Figure 5:**
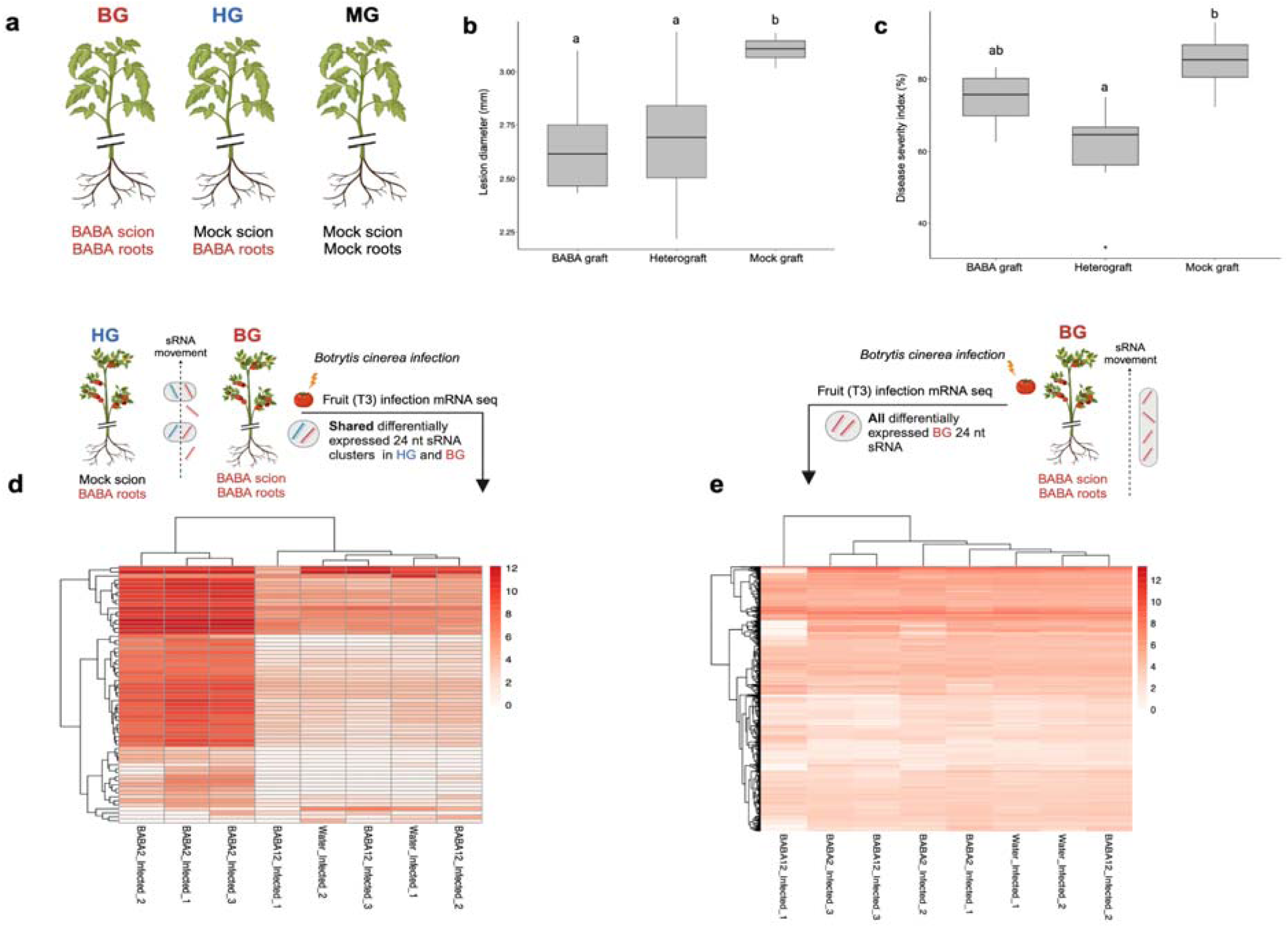
a) Experimental set up for grafting: from left to right BABA graft (BG) consisting of BABA scion and BABA treated roots scion, heterograft (HG) consisting of water scion and BABA roots and Mock graft (MG) consisting of Mock scion and Mock roots. b) Lesion diameter in tomato leaves infected with *B. cinerea* at 3 days post infection and c) disease severity index in ripe tomato fruit converted from the percentage of lesions in 6 disease categories at 6 days post infection (dpi) with *B. cinerea*. Different letters denote significant differences among treatment groups (one-way ANOVA, LSD post hoc test, p < 0.05, n = 3-8). Heatmap displaying normalised fragments per kilobase of transcript per million mapped reads (FPKM) value of genes containing d) upregulated differentially expressed 24 nt sRNA clusters conserved to HG (blue) and BG (red) phenotype captured in grey horizontal oval circles. FPKM values display expression level in fruit tissue of BABA2 infected, water infected and BABA12 *B. cinerea* infected plants. e) Upregulated differentially expressed 24 nt sRNA clusters in BG phenotype (red) captured in a grey vertical oval circle. FPKM values display expression level in fruit tissue of BABA2 infected, water infected and BABA12 *B. cinerea* infected plants.

Considering that the greatest changes in DNA methylation in our resistance phenotype were found in the CHH context and that this type of methylation is known to be mediated by RdDM, we tested whether the resistance phenotype in HGs is also associated with a change in accumulation of small RNAs (sRNAs). Scion leaf tissue was harvested 6 weeks after grafting and sRNA sequencing was conducted. Differentially expressed (DE) clusters were identified by comparing BG and HG with the MG clusters. In all cases, the majority of DE clusters were sRNAs 24 nt in size - (HG v MG: 47%, HG v BG: 92%, BG v MG: 85%). We observed a much larger number of DE clusters in the BG plants (17,323 up-regulated and 100,298 down-regulated) than the HG plants (10,150 up-regulated and 38,757 down-regulated). In contrast, HG vs BG contained 14,790 up-regulated DE clusters and 108,516 down-regulated DE clusters. This result indicates that the HG scions maintained an intermediate sRNA profile between MG and BG plants but are more similar to the former condition. This suggests that BABA treatment induced both graft transmissible and non-transmissible effects on sRNA accumulation.

To identify BABA-induced effects more directly linked to long-term resistance, we identified differentially expressed 24 nt clusters present in both resistant phenotypes (HG and BG) compared to control (MG), producing a total list of 6,010 clusters (Supp Table 2). From these, we identified 135 and 119 genes overlapping up-regulated and down-regulated 24 nt size clusters, respectively. To assess whether these sRNAs were associated with priming responses, the expression levels during *B. cinerea* infection of these overlapping genes were assessed by comparing their fragments-per-kilobase-million (FPKM) values across all treatment groups for infections at T1 and T3 (Fig 5d).

In the 135 genes coinciding with up-regulated sRNA clusters, we observed a clear trend of enhanced overexpression in response to infection in the BABA2 T3, compared to all other conditions (Fig 5d, Supp Fig 2a). For these genes, statistically significant enrichment was observed in the following biological process: ‘COPII-coated vesicle budding’ (103.4 fold enrichment, FDR p-value 0.025), ‘Aspartate family amino acid biosynthetic processes’ (74.4 fold enrichment, FDR p-value 0.025), ‘Vesicle budding from membrane’ (71.6 fold-enrichment, FDR p-value 0.025), and ‘Aspartate family amino acid metabolic processes’ (51.7 fold-enrichment 0.029 p-value FDR). Notably, such enhanced response to infection seen in BABA2 T3 is largely specific for these 135 genes, and it is not observed when all genes overlapping upregulated 24 nt clusters from the only BG condition are considered (2279 genes) (Fig 5e, Supp Fig 2b). Similarly, this effect was not observed for the BABA2 T3 119 genes associated with down-regulated clusters (Supp Fig 2c), or in the BABA2 T1 condition for both genes associated with up-regulated and down-regulated clusters (Supp Figs 2e-f). Therefore, our results suggest that mobile sRNAs accumulating in resistant naive tissue grafted on BABA-primed rootstock are specifically associated with a set of genes with an enhanced response to infection in tomato fruit expressing long-lasting induced resistance.

## Discussion

Here, we have demonstrated that plant age is a critical factor for the generation and maintenance of long-lasting induced resistance through chemical priming of defence. We have identified that BABA treatment of 2-week-old tomato seedlings (BABA2) leads to long-lasting postharvest and heritable resistance, and that this resistance phenotype is not observed if BABA treatment is conducted in 12-week-old plants (BABA12) (Fig 1b). These results suggest that young plants respond to BABA treatments differently than older plants. It is well known that due to the sessile nature of plants, lasting ‘memories’ can be created over the plant’s lifetime to facilitate survival and adaptation to environmental pressures^35^. This memory formation and maintenance has been demonstrated to occur via changes to epigenetic marks^36^. Moreover, significant epigenetic modifications in chromatin structure and DNA methylation are known to occur during development^37^. For example, two-day old Arabidopsis leaves have a heterochromatin content of 6% which increases to 14% at 2-weeks, thus increasing chromatin packing and impacting transcriptional activity^38,39^. We used this understanding as a basis to investigate whether the postharvest resistance observed in seedlings could be due to development-dependent differences in epigenetic profiles. Evidence of a dynamic epigenetic landscape throughout their lifetime is consistent with the idea that young plants could display greater epigenetic plasticity, thus facilitating robust memory formation. We therefore suggest that BABA seedling treatment leads to early foundational changes within the epigenome, facilitated by the young plant’s enhanced plasticity. In addition, these changes might result in pivotal effects in terms of capacity for subsequent epigenetic reprogramming and consequently, gene expression.

Interestingly, our DNA methylation analysis displayed negligible changes in the global methylation levels following BABA treatment at 2 or 12 weeks (Fig 3a), while similar previous experiments performed on the variety Money-Maker have found that BABA induces global DNA hypomethylation in the tomato variety Money-Maker^14^. This contrasting observation within the same plant species and priming elicitor could be due to the known differences among tomato varieties in perception of BABA^26^ and in the dynamic resource reallocation from growth to defence, which manifests mainly as a growth reduction^8,28^. We therefore speculate that the global changes in methylation levels in Money-Maker are associated with BABA-induced stress upon perception of the elicitor.

Despite the absence of global methylation changes found between treatments in our experiments (Fig 3a-c), DMR analysis unravelled clear differences between short-term and long-term priming responses. Specifically, we observed that early treated plants display a unique pattern of CHH hyper-DMRs and a higher number of DMRs months after the initial priming treatment, with BABA2 fruit having nearly double the number of CHH DMRs than BABA12 (Fig 3f). These observations could be directly associated to IR, as CHH methylation has been described to play a relevant role in response to stimuli and stress^40^ and has become recognised as an epigenetic modification marking the maintenance and expression of long-lasting priming^14,41^. Nonetheless, DMRs created by the BABA2 treatment are not maintained throughout the life of the plant. This supports the hypothesis that while BABA treatment has a pivotal influence on the future DNA methylation landscape, the early methylation response to BABA may not be directly associated with the establishment of long-term IR. Moreover, CHH methylation levels in leaf tissue double over 10 weeks from 7.3% to 13.5% (Fig 2c), indicating a highly dynamic profile of this epigenetic mark in tomato development. This agrees with various studies that have demonstrated that global methylation levels increase with chronological age in many plant species^42^. For example, Valledor demonstrated needle maturation of *Pinus radiata* is associated with an increase in global DNA methylation of ∼3% over 12 months^43^. Taking this observation into consideration, we can argue that the lower global CHH level in young seedlings is associated with a greater plasticity in young plants and facilitates a strong priming imprinting following BABA treatment.

The observation that DMRs associated to BABA2 are not maintained from seedlings to fruiting (Fig 3h) further supports the hypothesis on the importance of foundational changes at early stages of development and tissues that experience early BABA exposure. Considering that tissues present early during development are influencing responses in fruit that were not present at the time of BABA2 treatment, this brings the question of whether the long-lasting resistance is due to a systemic signal. We therefore combined the use of grafting and BABA soil drench treatments. Our experiments demonstrate that long-lasting resistance is transmissible across graft junctions to untreated naive scions (Fig 5b-c). The absence of long term-resistance observed in BABA12 plants exclude that this primed state is due to residual BABA transported to and accumulated in fruit tissue^16^.

Our efforts then focused on the characterisation of this mobile signal by investigating the accumulation of sRNAs, which are known to be mobile and able to transmit traits across a graft junction^44^. Previous grafting experiments performed with transgenic plants have demonstrated that 24 nt sRNAs lead to epigenetic changes in recipient scion tissue^45^. Importantly, the RdDM pathway has been frequently associated with defence against biotic stresses and with the expression and maintenance of long-lasting priming. For example, the double mutant *drm1drm2* displays enhanced susceptibility against the necrotrophic pathogen *P. cucumerina*^40^. Furthermore, the mutant *nrpe1* was shown to display enhanced resistance against *Pseudomonas syringae* DC3000, with enhanced SA and reduced JA-mediated defences respectively^40^, but fails to generate transgenerational resistance^46^. Therefore, one could speculate that the RdDM pathway plays a role in the establishment and maintenance of long-lasting priming in tomato by impacting mobile sRNAs that alter methylation in naive tissue.

To identify siRNAs potentially driving the resistance observed in our experiments, sRNA sequencing was conducted on leaf tissue from grafted plants. Interestingly, while BABA treated plants (BG) display a change in the accumulation of many sRNA clusters, only a small portion are maintained in grafted scions (HG) and are therefore associated with long-lasting resistance. This result suggests that most BABA induced effects on sRNA accumulation are not involved in graft-transmissible IR. Conserved differentially expressed 24 nt sRNA clusters were identified from our resistant plants (BG and HG). The 135 genes associated with up-regulated clusters displayed a clear pattern of enhanced expression during fruit infections in resistant BABA2 plants (Fig 5d). Enhanced expression of these cluster associated genes was not observed in BABA12 or water plants. Moreover, the same expression pattern was not observed in leaf tissue or across our down-regulated cluster associated genes, (Supp Fig 2c-f). Whereas this could be due to the dynamic nature of transcriptomic responses during pathogen infection and the limitations of collecting single timepoints^47^, our results suggest that early BABA treatment generates a systemic signal that leads to dynamic changes in the priming response in the fruit.

Among the genes that we found functionally associated with sRNA clusters and priming in our experiment, significant enrichment was observed in pathways including ‘COPII-coated vesicle budding’ and ‘Aspartate family amino acid biosynthetic process’. The latter is particularly intriguing, since priming by BABA in Arabidopsis has been directly associated with aspartic acid-related processes, following identification of the BABA receptor as the aspartyl-tRNA synthetase *IBI1*^26^. In addition, an asparagine synthetase was also found in these T3 primed genes, providing further evidence of a role of amino acid processes related to aspartic acid. Interestingly, this asparagine synthetase has been shown to be important in defence against microbial pathogens and is mainly found in root tissue^48,49^, where BABA is applied and taken up.

Our experiments were part of an approach to identify epigenetic markers to develop protection strategies against *B. cinerea* in tomato. The results have the potential to impact the horticulture industry, as they guide it towards priming applications and imprinting of resistance phenotypes at early stages of cultivation, thereby reducing associated commercial costs. Moreover, this work has led to the discovery that long-lasting BABA-IR is transmissible through graft junctions, and we have identified a group of sRNAs with a potential role in DNA methylation-dependent maintenance of long-lasting priming. Considering the relevance of grafting to the horticulture industry due to enhanced plant vigour^44,50^, our results could be exploited into commercial settings, for both enhanced growth and protection against diseases pre- and postharvest.

## Materials and Methods

### Plant materials and growth conditions

Seeds of tomato cv Micro Tom were placed in Petri dishes containing wetted tissue paper and maintained at 28°C in the dark for 3-6 days to stimulate homogeneous germination. Germinated seeds were planted in individual pot propagators containing Scott’s M3 compostand cultivated under standard tomato growth conditions (16h/8h day/night cycle; 25°C/20°C).

### BABA treatments and sample collection

At two-weeks, 1/3 of plants were soil drenched with 5 mM BABA (catalogue number A4420-7, Sigma Aldrich) to provide a final concentration of 0.5 mM in the soil. All other plants were treated with the same volume of water. One week after BABA treatment, seedlings were removed from soil, roots were gently washed, and seedlings were replanted in new compost. At twelve weeks, 1/3 of plants were soil treated to a final concentration of 0.5 mM BABA. The remaining 1/3 of plants were left as water controls. One-week after each respective BABA treatment, two leaves from four individual plants were collected and stored in liquid nitrogen. Plants were left to grow and produce red fruit. The first three fruit on all plants were marked.

### Relative Growth Rate analysis

Plant height was measured before BABA treatment at 2 weeks (t1) and one week after BABA treatment (t2), as previously described^28^. Relative growth rate (RGR) was calculated using the formula = LN(height-t2)-LN(height-t1)/t2-t1^51^.

### Botrytis cinerea infections

Infections of leaves with *Botrytis cinerea* (R16) were performed as previously described^28^, with minor modifications. Briefly, each tomato leaf was infected with droplets of 5 μL inoculum containing 5×10^5^ spores/ml. Leaves were incubated in the dark at 100% humidity and 20°C. Disease was scored by measuring lesion diameters using an electronic calliper at 2 days post infection (dpi). Infection of fruit was performed entirely as described in Wilkinson et al., 2018^16^. Disease was scored by classifying fruit in different categories of infection and distributions were represented as disease severity index as previously described by Wilkinson et al., 2018^16^. Statistical analysis was done in R (version 1.4.1717) and as previously described in Luna et al., 2016 and Wilkinson et al., 2018^16,28^.

### Nucleic acid extractions

At 15-weeks the first fruit was collected from each plant and stored in liquid nitrogen for DNA extraction for methylation analysis. DNA was extracted from 100 mg of ground frozen leaf and fruit tissue using Qiagen DNeasy Plant Mini Kit (Qiagen) following the manufacturer’s instructions and resuspended in 30 μL H_2_O. For transcriptomic analysis, leaf and fruit tissue was collected at 1 and 2 dpi respectively. RNA for all experiments was extracted from 100 mg of ground frozen tissue using TRIzol (Thermo Fisher; catalogue number 15596026) following manufacturer’s specifications.

### Library preparation and sequencing mRNA data

mRNA sequencing analysis was conducted on RNA extracted from fruit tissue 48 hour post infection (hpi) with *B. cinerea*. Library preparation and sequencing was conducted by Novogene using NovaSeq 6000 PE150. 20 million paired reads were generated per sample. A total of 767,502,726 clean reads were generated across 17 samples, with an average of 45,147,219 clean reads per sample. An average of 92.6% of nucleotides per sample had a Phred quality score of > 30. The quality of samples was assessed using FAST QC (0.11.5-Java-1.8.0_74). Adapters were removed from samples using Trimmomatic (version 0.39)^52^. Reads were aligned to the tomato genome (ITAG4.0) using Hisat2 (version 2.2.1)^53^. Samtools (version 1.12)^54^ was used to sort and index Sam files into Bam format. Read counts were generated using Htseq (version 0.13.5)^55^. All sequencing information is summarised in Supp Table 3.

### Statistical analysis of sequence count files

Count files were analysed in R using deseq2 (version 1.34.0)^56^. Genes with total read counts of under 10 were removed. A variance stabilising transformation (VST) was applied for normalisation, principal component analysis (PCA) plots were generated using deseq2 and ggplot2, and hierarchical clustering was conducted for samples using Euclidean distances. Differentially expressed genes (DEGs) involved in infection were identified for infected plants of all three conditions (BABA2, Water, BABA12). DEGs were identified between BABA2 and water, BABA12 and water plants to identify the effect of early and late BABA treatments on the transcriptome without infection.

### Library preparation and whole genome bisulfite sequencing processing

All library preparation and sequencing was conducted by Novogene using NovaSeq 6000 PE150. After quality control, positive control DNAs were added into the DNAs which were then fragmented into 200-400 bp using Covaris S220 and bisulfite treated (Accel-NGS Methyl-Seq DNA Library Kit for illumina, Swift). Ligation of methylation sequencing adapters, size selection and PCR amplification was then performed before illumina sequencing for 20 million paired reads per sample with 30 x sequencing depth. An average of 90.2% of nucleotides per sample had a Phred quality score of > 30. A total of 1.09938788 × 10^9^ reads were generated across all 24 samples, with an average of 45.8 million clean reads per sample. The tomato genome version 4.0 was downloaded from solgenomics.net/organism/Solanum_lycopersicum/genome. The genome was merged with the chloroplast sequence (NC_007898.3) to calculate the bisulfite conversion rate. The reference genome was bisulfite converted using Bismark (v0.16.3_bowtie2)^57^. Quality of samples was assessed using FAST QC (0.11.5-Java-1.8.0_74). Adapters were removed from samples using Trimmomatic (version 0.39)^52^ using the following parameters “LEADING:30 TRAILING:30 SLIDINGWINDOW:4:15 MINLEN:16”. Reads were aligned to the SL4.0 tomato genome using Bismark^57^ “Bismark -N 1 -L 20 -p 4 -X 1000 -score_min L,0,-0.6” which uses Bowtie for mapping. The average mapping efficiency was 80% with a minimum and maximum value of 77% and 82.2% respectively. Samples were de-duplicated using Samtools (/1.8-iomkl-2018a). CX files were created with Bismark using: “bismark_methylation_extractor –comprehensive --multicore 8 --bedGraph --CX --buffer_size 10G”. The bisulfite conversion rate was calculated for each sample using the chloroplast sequence; conversion rates varied from 98.41-99.66%. An estimated conversion rate was applied following Catoni and Zabet, (2021)^58^ in order to account for non-converted DNA. All sequencing information is summarised in Supp Table 3.

### Analysis of DNA methylation

Differentially methylated regions (DMRs) were identified starting from the Bismark CX files using a previously established pipeline^58^. Briefly, we used the package *DMRcaller* (version 1.26.0)^59^ to identify DMRs in each cytosine context (CG, CHG and CHH) in all paired combinations of our samples, including three different timepoints: T1 (3 weeks) Water and BABA2 leaf tissue one week post BABA soil drench; T2 (13 weeks) Water, BABA2 and BABA12 leaf tissue one week post BABA soil drench; and T3 (15 weeks) Ripe red fruit in water, BABA2 and BABA12. The following parameters were used: function *computeDMRsReplicates*, the ‘bins’ method was used with a bin size of 200 bp, methylationDiff of 0.2, minCytosinesCount of 2, minReadsPerCytosine of 4, and a minimum *p-*value of 0.05. To identify DMRs unique to the ‘BABA2’ treatment group, overlap was conducted between BABA2 and BABA12 DMRs, with only BABA2 DMRs used for further downstream analysis. To identify genomic features, the ITAG4.1 tomato reference genome was used. Repeat Modeler was used to identify repeats^60^. The R package *GenomicRanges* (version 1.46.1)^61^ was used to assess overlap between DMRs and genomic features.

### GO enrichment analysis

GO enrichment analysis was performed on differentially expressed genes and genes overlapping differentially methylated regions (+/- 2 kb). Analysis was performed by using the online tool ShinyGO^62^ with Fisher’s exact test, false discovery rate correction and a *p* value cut-off of 0.05.

### Grafting

To assess if BABA-IR is transmissible through grafting, two-week old Micro Tom plants were treated to a final concentration of 0.5 mM BABA via soil drench treatment or with water. 3 days after soil treatments, grafting was conducted on seedlings using the cleft method^63^. The following graft combinations were used with a minimum of 4 biological replicates per treatment: BABA graft (BG) BABA treated root stock + BABA treated scion; Hetero graft (HG) BABA treated root stock + water treated scion; and Mock graft (MG) water treated root stock + water treated scion.

### Library preparation and sequencing sRNA data

sRNA sequencing analysis was conducted on RNA extracted from scion leaf tissue from BG, HG and MG plants 6 weeks post grafting. Library preparation and sequencing was conducted by Novogene using NovaSeq SE50. 20 million paired reads were generated per sample. A total of 302,217,035 clean reads were generated across 12 samples, with an average of 25,184,752 clean reads per sample. An average of 97.5% of nucleotides per sample had a Phred quality score of > 30. All sequencing information is summarised in Supp Table 3.

### Analysis of sRNAs

Quality of data was assessed using FAST QC. Samples were trimmed using Trimmomatic (version 0.39) to remove adapter sequences and obtain reads between 16-30 bp in length^52^. Trimmed reads were uniquely mapped to the reference genome using Shortstack (version 3.8.5)^64^. Clusters were identified as 20-24 nt in size using the *mobileRNA* package (https://github.com/KJeynesCupper/mobileRNA). For each replicate the consensus cluster size was identified and non-dicer-derived clusters were excluded from the analysis. Differentially expressed clusters were identified between HG and MG, BG and MG and HG and BG using the DEseq2 method. Location of DE clusters was found using *GenomicRanges* (version 1.46.1)^61^ by overlapping clusters (+/- 1kb) with known genomic features using a log threshold change of 1 or −1 for up and downregulated, respectively; a false discovery rate (FDR) threshold of 0.05 was applied to control significance.

## Acknowledgements

We thank Dr S. W. Wilkinson for extremely valuable discussions. This work was funded by the BBSRC Future Leader Fellowship BB/P00556X/1 and BB/P00556X/2 to E.L., and the Horticultural Quality and Food Loss Network pump-priming grant WXA3189N/P16188/UoB_Luna-Diez to E.L., M.C., M.R.R. and K.S. This work was possible thanks to the BBSRC and University of Birmingham Midlands Integrative Biosciences Training Partnership (MIBTP) BB/M01116X/1 iCASE studentship to K.S., which is supported by the Agri-tech company Saturn Bioponics. Work by M.C. is funded by the BBSRC responsive mode grant BB/W008866/1. K.J-C. is supported by the MIBTP grant BB/T00746X/1.

## Author contributions

E.L., M.C. and M.R.R. conceived the study and obtained core funding. E.L., K.S. and M.C. designed the experimental work pipeline. K.S. conducted experiments and gathered data. E.L, K.S., M.C. and M.R.R designed the data analysis pipeline. K.S. performed data analysis with the guidance of K.J-C on the analysis of the sRNA data. K.S. and E.L. performed statistical analyses and data interpretation with the support of M.C., M.R.R. and K.J-C. K.S. and E.L. drafted the first version of the article. K.S., E.L., M.C. and M.R.R. wrote the submitted version of the article.

## Source Data

Sequencing data has been deposited in Gene Expression Omnibus under the accession number (*awaiting allocation*).

