## Supplementary material for "Developmentally regulated generation of a systemic signal for long-lasting defence priming in tomato": Supp Fig 1

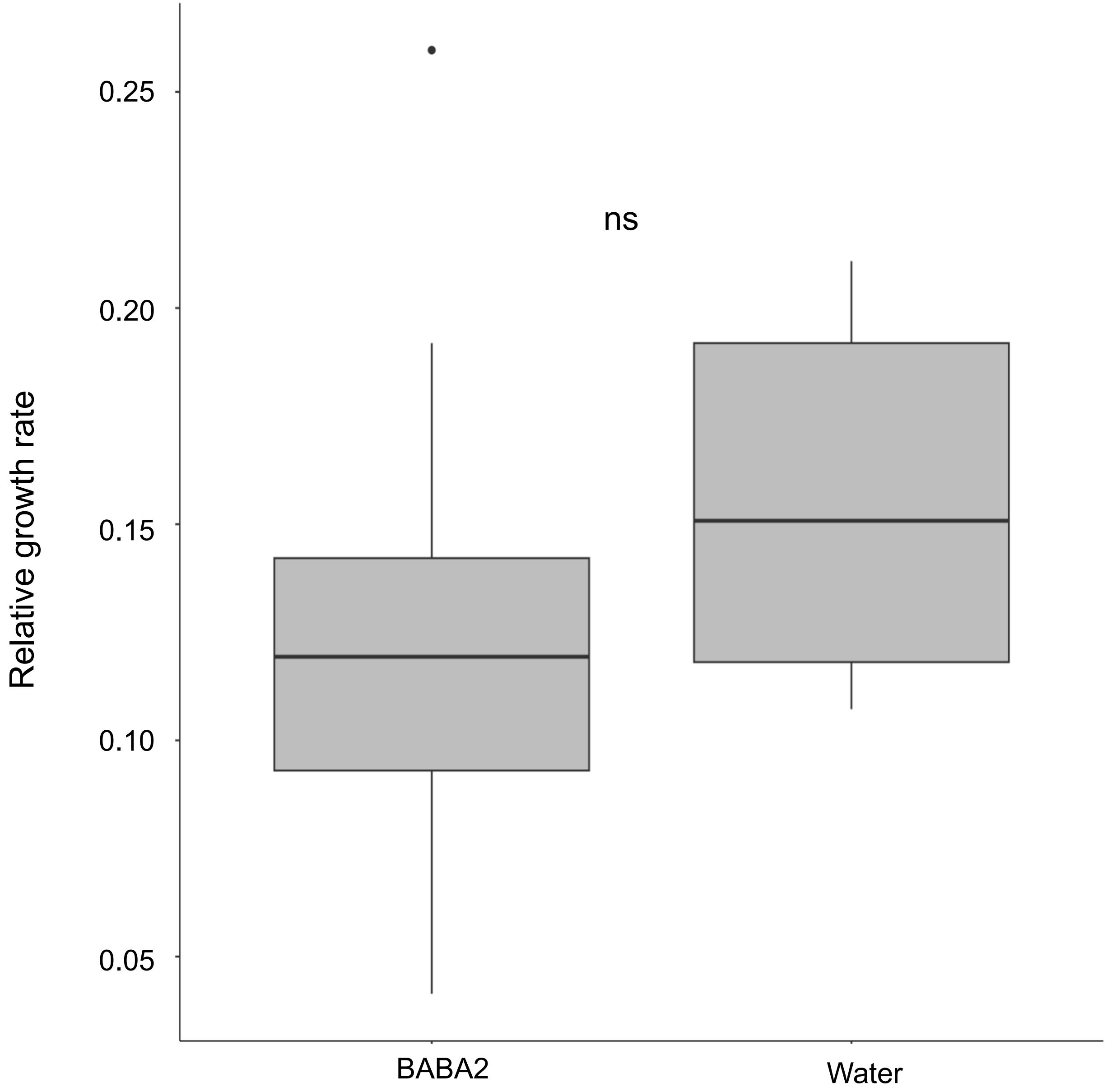


**Fig S1** – Average relative growth rates (in cm per cm per week; ± standard error of the mean [SEM]) of plants between 2 and 3 weeks after 0.5 mM BABA treatment. Ns denotes no significant differences among treatment groups (*t* test, $p = 0.05$; $n = 8\text{-}10$)
