## Supplementary material for "Developmentally regulated generation of a systemic signal for long-lasting defence priming in tomato": Supp Fig 2

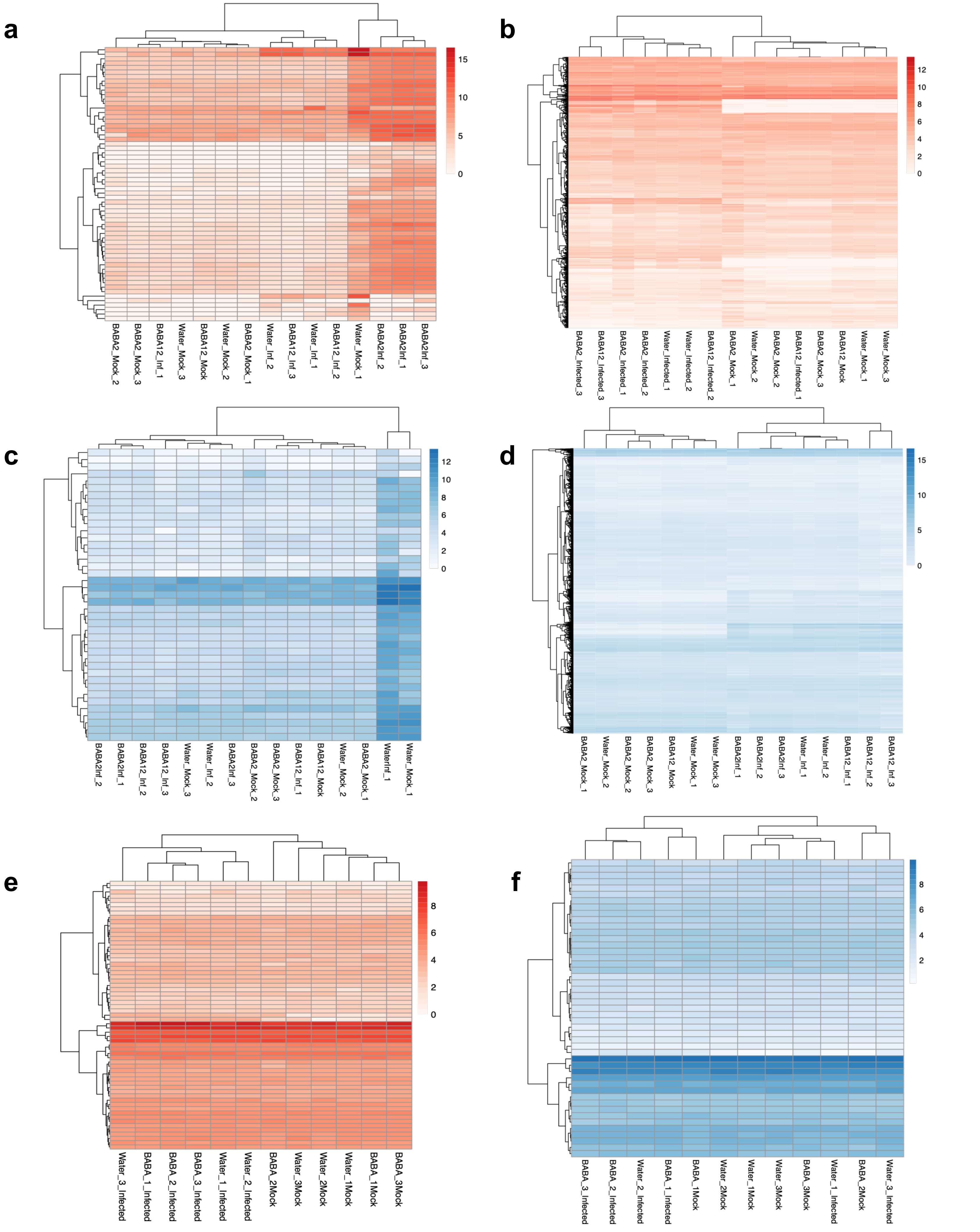


**Fig S2:** Heatmap displaying normalised fragments per kilobase of transcript per million mapped reads (FPKM) value of genes containing a) upregulated differentially expressed 24 nt sRNA clusters conserved to HG and BG phenotype. FPKM values display expression level in fruit tissue of mock and *B.* cinerea BABA2, Water and BABA12 plants. b) Upregulated differentially expressed 24 nt sRNA clusters in BG phenotype. FPKM values display expression level in fruit tissue of mock and *B. cinerea* infected BABA2, Water and BABA12 plants. c) downregulated differentially expressed 24 nt sRNA clusters conserved to HG and BG phenotype. FPKM values display expression level in fruit tissue of mock and *B.* cinerea BABA2, Water and BABA12 plants. d) Downregulated differentially expressed 24 nt sRNA clusters in BG phenotype. FPKM values display expression level in fruit tissue of mock and *B. cinerea* infected BABA2, Water and BABA12 plants. c) upregulated differentially expressed 24 nt sRNA clusters conserved to HG and BG phenotype. FPKM values display expression level in leaf tissue of mock and*B.* cinerea BABA2, Water and BABA12 plants (T1). f) Downregulated differentially expressed 24 nt sRNA clusters in BG phenotype. FPKM values display expression level in leaf tissue of mock and *B. cinerea* infected BABA2, Water and BABA12 plants (T1).
